# Aggressive high-grade NF2 mutant meningiomas downregulate oncogenic YAP signaling via the upregulation of VGLL4 and FAT3/4

**DOI:** 10.1101/2024.05.30.596719

**Authors:** Abigail G Parrish, Sonali Arora, H. Nayanga Thirimanne, Dmytro Rudoy, Sebastian Schmid, Philipp Sievers, Felix Sahm, Eric C Holland, Frank Szulzewsky

## Abstract

Meningiomas are the most common primary brain tumors in adults. Although generally benign, a subset of meningiomas is of higher grade, shows aggressive growth behavior and recurs even after multiple surgeries. Around half of all meningiomas harbor inactivating mutations in NF2. While benign low-grade NF2 mutant meningiomas exhibit few genetic events in addition to NF2 inactivation, aggressive high-grade NF2 mutant meningiomas frequently harbor a highly aberrant genome. We and others have previously shown that NF2 inactivation leads to YAP1 activation and that YAP1 acts as the pivotal oncogenic driver in benign NF2 mutant meningiomas. Using bulk and single-cell RNA-Seq data from a large cohort of human meningiomas, we show that aggressive NF2 mutant meningiomas harbor decreased levels YAP1 activity compared to their benign counterparts. Decreased expression levels of YAP target genes are significantly associated with an increased risk of recurrence. We then identify the increased expression of the YAP1 competitor VGLL4 as well as the YAP1 upstream regulators FAT3/4 as a potential mechanism for the downregulation of YAP activity in aggressive NF2 mutant meningiomas. High expression of these genes is significantly associated with an increased risk of recurrence. In vitro, overexpression of VGLL4 resulted in the downregulation of YAP activity in benign NF2 mutant meningioma cells, confirming the direct link between VGLL4 expression and decreased levels of YAP activity observed in aggressive NF2 mutant meningiomas. Our results shed new insight on the biology of benign and aggressive NF2 mutant meningiomas and may have important implications for the efficacy of therapies targeting oncogenic YAP1 activity in NF2 mutant meningiomas.

## Introduction

Meningiomas are the most common primary brain tumors in adults ^1^. Although only a subset of meningiomas is of higher-grade and show aggressive growth behavior, atypical and anaplastic meningiomas frequently invade into the brain and tend to recur even after multiple rounds of surgery, chemo-, and radiation therapy. Around half of all meningiomas exhibit functional loss of the *NF2* gene, either due to loss of chromosome 22, inactivating point mutations, or gene fusions. Aggressive meningiomas occur in all molecular subgroups but are enriched in NF2 mutant meningiomas. While benign NF2 mutant meningiomas usually only exhibit functional NF2 loss and rarely harbor additional recurrent mutations or chromosomal aberrations, aggressive NF2 mutant meningiomas generally harbor a more aberrant genome with several recurrent chromosomal gains and losses in addition to functional NF2 inactivation/chromosome 22 loss, such as losses of chromosomes 1p, 4, 6, 10, and 14 ^2,3^.

NF2 (encoding for the protein Merlin) is a potent tumor suppressor. In addition to suppressive functions in the Ras-Raf-ERK and PI3K-AKT-mTOR-S6 pathways ^4,5^, NF2 predominantly functions as an upstream regulator of the Hippo Signaling pathway that translates mechanical stimuli into transcriptional signals ^6-8^. NF2, in a complex with Expanded (FRMD6) and Kibra (WWC1), activates a core cascade of serine-threonine kinases (STK3/4, LATS1/2) that ultimately phosphorylate the transcriptional co-activator YAP1 (and its paralog TAZ) at five key serine residues (most importantly S127 and S397) ^9,10^. Phosphorylation of YAP1 at these residues results in its functional inhibition by both cytoplasmic retention and proteasomal degradation. Several studies have detected increased YAP activity in NF2 mutant meningioma tumors and/or cell lines ^11,12^. Recently, we have shown that the expression of a non-regulatable YAP1 variant (S127/397A-YAP1) induces very similar transcriptional changes compared to NF2 loss in human neural stem cells and that the expression of the same construct in Nestin-positive cells in the meninges of *Cdkn2a* null mice induces the formation of meningioma-like tumors that resemble human NF2 mutant meningiomas by histomorphology and gene expression ^13^. In addition, recurrent YAP1 gene fusions have been identified in a subset of pediatric NF2 wild type meningiomas ^14^ and other CNS and non-CNS tumors ^6,15^. The oncogenic functions of these YAP1 fusions fundamentally rely on their ability to exert de-regulated YAP activity and YAP1 fusion-positive human meningiomas resemble NF2 mutant meningiomas by both gene expression and DNA methylation-based classification ^13-15^, further linking oncogenic de-regulated YAP activity to NF2 loss and meningioma-genesis. In addition to the canonical Hippo Signaling pathway working through NF2, additional upstream Hippo regulators have been identified that can either act through the canonical STK-LATS axis or via non-canonical Hippo signaling. Gigantic cadherin proteins FAT and Dachsous positively regulate Hippo signaling and inhibit Yorkie (the YAP1 ortholog in Drosophila). In Drosophila, FAT can activate the Hippo pathway independently of NF2, and, while the exact mechanism is not conserved in mammals, mammalian FAT family proteins (FAT1-4) have been implicated in the regulation of YAP1 ^16-19^. Loss or functional inactivation of FAT proteins has also been observed in benign NF2 wild type spinal meningiomas, suggesting tumor suppressive activity of FAT proteins in benign meningioma ^20^.

YAP1 is a transcriptional co-activator that predominantly functions through its interaction with the TEAD family of transcription factors ^21-23^. Ablation of the YAP1-TEAD interaction, either through mutagenesis or pharmacological inhibition, results in functional YAP inactivation. Ablation of YAP1-TEAD interaction by S94A mutation of YAP1 resulted in the inability to induce the gene expression changes and growth promoting effects elicited by wild type YAP1 ^23^. Similarly, pharmacological inhibition of the YAP1-TEAD interaction with small molecule inhibitors can inhibit the expression of YAP1 downstream targets (such as CYR61/CCN1 and CTGF/CCN2) as well as the growth of YAP1-driven cells. Several of these inhibitors are currently under investigation in clinical trials ^15,24,25^. Vestigial-like family members 1-4 (VGLL1-4) are transcriptional cofactors that compete with YAP1 for TEAD binding ^26-28^. VGLL1-3 each contain one Tondu (TDU) domain important for the interaction with TEADs, whereas VGLL4 contains two conserved TDUs that mediate the VGLL4-TEAD interaction. Except for their TDU domains, these proteins bear no sequence similarity, suggesting that VGLL1-3 and VGLL4 might have distinct molecular functions ^29,30^. Furthermore, while several studies have identified oncogenic functions of VGLL1-3 in different cancer types ^27,31-34^, VGLL4 has so far only been described as a tumor suppressor and reduced VGLL4 expression levels have been detected in several cancers ^35-38^. However, the exact functions of VGLL4 in tumorigenesis or progression remain unclear.

Historically, these tumors have been classified into three WHO grades, based on established histologic criteria. However, the histological grade frequently does not accurately predict tumor growth, behavior, and recurrence, highlighting the need for additional classifications based on genetic, epigenetic, and transcriptional markers. Recent large-scale sequencing efforts have utilized several approaches, including DNA methylation-based sequencing, whole exome sequencing, and bulk RNA sequencing to classify meningiomas beyond the classic WHO grading system ^2,3,39,40^. To gain a deeper insight into which pathways are activated in the different meningioma subclasses and grades, we have recently collected RNA Seq data of around 1300 human meningiomas and performed UMAP analysis to identify clusters that correlate with clinically relevant subgroups. We observed that RNA Seq analysis was able to predict tumor grade, recurrence, and functional NF2 status, and observed significant overlaps with clusters derived from DNA methylation-based and whole exome sequencing, suggesting that RNA Seq-based classification yields compatible results ^41^.

Several open questions remain about the pathobiology of both benign and aggressive NF2 mutant meningiomas. In this study, we utilize our collection of RNA-Seq data from 1300 human meningioma samples as well as additional human meningioma single-cell sequencing data to investigate the levels of YAP1 activity in the different meningioma subclusters. We find that aggressive NF2 mutant meningiomas exhibit decreased levels of YAP1 activity, the putative oncogenic driver in benign NF2 mutant meningiomas. We then identify elevated expression of the YAP1 antagonist VGLL4 in a subset of aggressive NF2 mutant meningiomas as one possible route of functional YAP1 inhibition. Lastly, using an in vitro cell culture model we show that overexpression of VGLL4 in benign NF2 mutant meningioma cells leads to the down-regulation of canonical YAP1 target genes and growth inhibition.

## Results

### Aggressive NF2 mutant meningiomas harbor reduced levels of YAP activity compared to benign NF2 mutant meningiomas

We have previously collected a large cohort of 1298 bulk RNA-Seq samples of human Meningiomas and have applied Uniform Manifold Approximation and Projection (UMAP), a dimensionality reduction method, on batch-corrected, normalized transcript counts to create a reference UMAP. DBSCAN (density-based spatial clustering of applications with noise) was then used to delineate regional distinct clusters that corresponded with metadata (such as time to recurrence, WHO grade) was able to separate NF2 mutant meningiomas into a benign and an aggressive cluster .

To explore the levels of YAP activity in different subtypes of meningioma (NF2 wild type, benign and aggressive NF2 mutant), we analyzed the expression of NF2, YAP1, and the YAP1 paralogue TAZ/WWTR1 in our cohort of 1298 bulk RNA-Seq samples from human meningiomas, including both NF2 mutant and wild type tumors. We subdivided NF2 mutant meningiomas into benign (n = 290) and aggressive (n = 476) tumors based on the previously established clusters (Figure 1A, Suppl. Figure S1A-B) ^41^. Since the activity of YAP1 is largely regulated at the protein level and not at RNA level, we also investigated the expression of several YAP1 target genes (CYR61/CCN1, CTGF/CCN2, AMOT, AMOTL2, ANKRD1).

**Figure 1:**
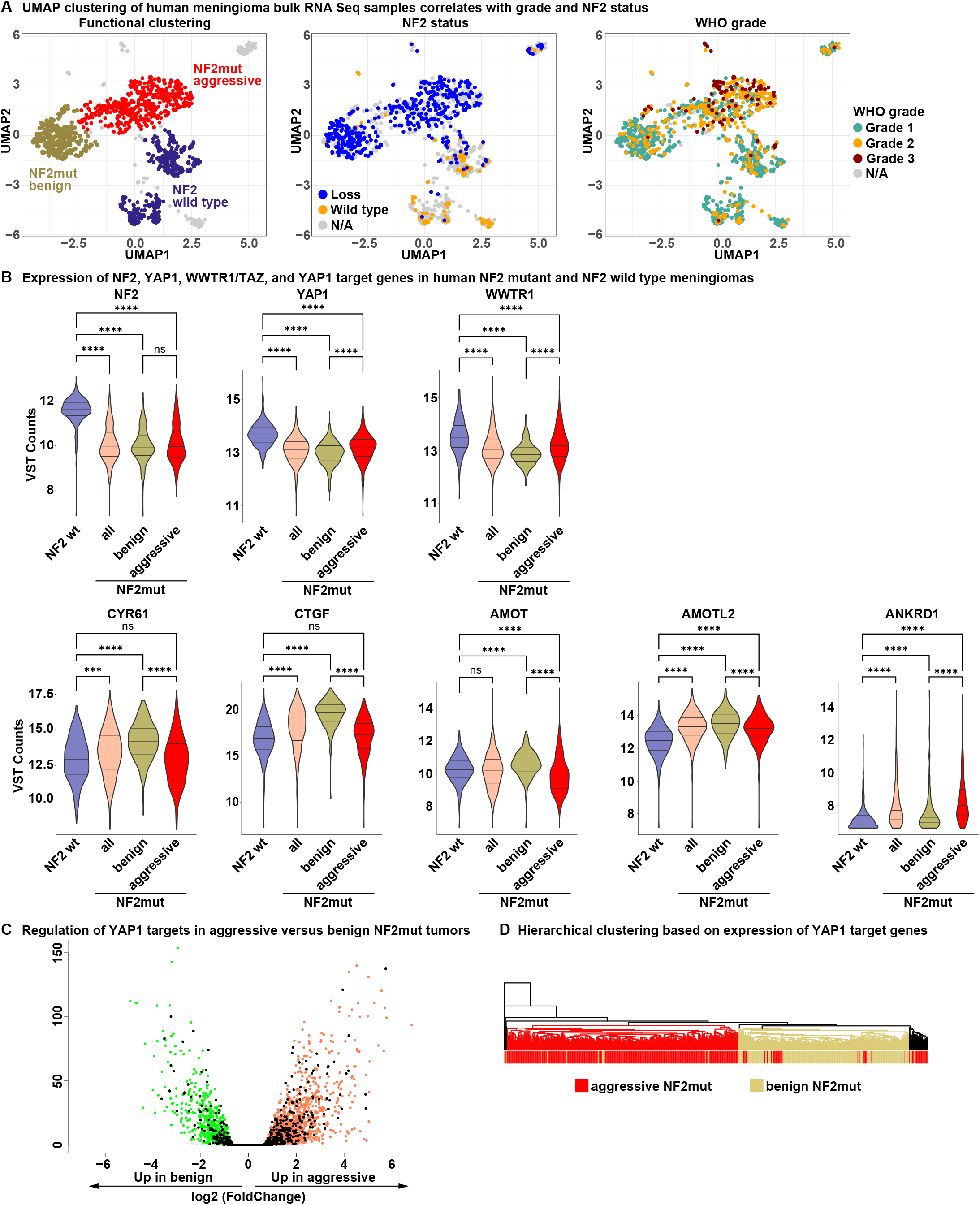
Aggressive NF2 mutant meningiomas display decreased levels of YAP activity. Reference UMAPs showing clustering of human meningiomas based on bulk RNA-Seq data. Samples are colored by cluster (benign NF2 mutant, aggressive NF2 mutant, NF2 wild type), NF2 status, or WHO grade. B) Expression of Hippo pathway genes and YAP1 target genes in bulk RNA-Seq data of different clusters of human meningiomas. C) Expression of YAP target genes (black dots) in benign versus aggressive human NF2 mutant meningiomas. D) Hierarchical clustering of human meningioma bulk RNA-Seq samples based on the expression of 2212 YAP target genes is able to separate benign and aggressive NF2 mutant meningiomas. Statistical analysis was done with One-way ANOVA (B). (***) P _≤_ 0.001; (****) P _≤_ 0.0001.

The expression of NF2, YAP1, and TAZ were all significantly decreased in NF2 mutant compared to NF2 wild type tumors. By contrast, the expression of the YAP1 target genes CTGF, CYR61, AMOTL2, and ANKRD1 was significantly higher in NF2 mutant tumors compared to NF2 wild type meningiomas (Figure 1B).

When comparing aggressive to benign NF2 mutant tumors, we detected no significant difference in the expression of NF2 between benign and aggressive NF2 mutant meningiomas, whereas the expression of YAP1 and TAZ was significantly higher in aggressive NF2 mutant tumors. By contrast, we detected significantly decreased expression levels of the YAP1 target genes CTGF (3.2-fold), CYR61 (2-fold), AMOT (1.7-fold), and AMOTL2 (1.5-fold) in aggressive compared to benign NF2 mutant meningiomas (Figure 1B, Suppl. Figure S1C). Of note, the YAP1 target gene ANKRD1 was expressed at significantly higher levels in aggressive versus benign NF2 mutant tumors (1.8-fold).

To analyze the differences in YAP activity between benign versus aggressive meningiomas on a broader scale, we calculated the differentially-expressed genes between benign and aggressive meningiomas (Fold change _≥_ 1.5, p-adjusted _≤_ 0.05). We observed 1836 up-regulated and 843 down-regulated genes in aggressive NF2 mutant meningiomas (compared to benign NF2 mutant tumors). We compared these differentially expressed genes (DEGs) to a dataset of YAP1 target genes ^42^ and observed that 17.3 percent (382 out of 2212 genes) of YAP1 target genes were significantly up-or downregulated between benign and aggressive NF2 mutant meningiomas (p = 5.34 x 10^-27^) (Figure 1C). Unsupervised hierarchical clustering based on the expression of these 2212 YAP1 target genes was also able to separate benign from aggressive NF2 mutant meningiomas (Figure 1D).

Taken together, our data suggests that aggressive NF2 mutant meningiomas harbor different levels of YAP activity and downregulate several YAP target genes compared to benign NF2 mutant meningiomas.

### Decreased YAP activity in aggressive meningiomas is also observed at the single cell level

Since benign NF2 mutant meningiomas exhibit increased immune cell infiltration compared to their aggressive counterparts ^43^, we used single-cell RNA sequencing data from 26 human meningiomas to assess the expression of YAP1 target genes in the tumor cells proper at the single-cell level ^43^ (Suppl. Table S1A). Samples were classified based on the Heidelberg methylation classifier ^3^ into either ben-1 (7 samples), ben-2 (3 samples), int-A (5 samples), int-B (2 samples), or mal (9 samples) subtypes. We excluded immune cells based on the expression of 49 marker genes (Suppl. Table S1B).

As observed in the bulk RNA-Seq samples from human meningiomas, we detected significantly decreased expression levels of several YAP1 target genes (CTGF, CYR61, ANXA3) in ben-1 NF2 mutant tumor cells compared to more aggressive NF2 mutant tumor cells, confirming that the down-regulation of YAP activity in aggressive meningiomas observed in bulk RNA sequencing samples is intrinsic to the tumor cells and not caused by contamination with microenvironment and immune cells (Figure 2A-B). Both CTGF and CYR61 were significantly downregulated in tumor cells of all the higher-grade/aggressive subtypes (NF2 mutant mal-subtype (CTGF: 2.48-fold decrease, padj = 1 x 10^-115^; CYR61: 3.32-fold decrease, padj = 4.5 x 10^-41^), int-B subtype (CTGF: 6.59-fold decrease, padj = 2.9 x 10^-162^; CYR61: 4.63-fold decrease, padj = 5.15 x 10^-40^), and int-A (CTGF: 3.18-fold decrease, padj = 1 x 10^-85^; CYR61: 2.77-fold decrease, padj = 1.33 x 10^-18^)) compared to samples of the ben-1 subtype.

**Figure 2:**
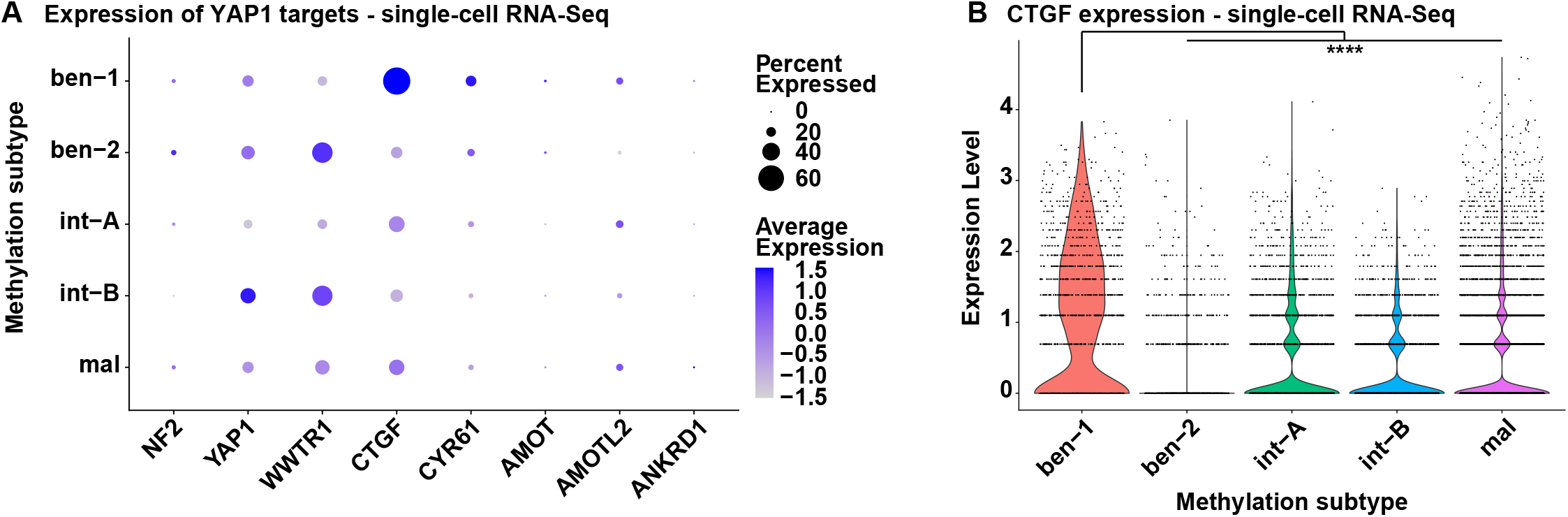
Expression of YAP target genes is decreased in aggressive type (int-A, -B, mal) human meningioma tumor cells on single cell level. A) Expression of Hippo Pathway effectors and YAP1 target genes in single-cell RNA-Seq data of human meningioma tumor cells of different methylation subtype tumor samples. B) Expression of the YAP1 target gene CTGF in single-cell RNA-Seq data of human meningioma tumor cells of different methylation subtype tumor samples. Statistical analysis was done with One-way ANOVA (B). (****) P _≤_ 0.0001.

Taken together, our results suggest that the decreased YAP activity in aggressive NF2 mutant meningiomas observed in bulk RNA sequencing data is also found in single-cell sequencing data and intrinsic to the tumor cells.

### Low expression of YAP1 targets in NF2 mutant meningiomas is associated with shorter time to recurrence

NF2 mutant meningiomas located in the benign and aggressive UMAP clusters are associated with a significantly different risk of recurrence (Figure 3A). We then assessed if the level of YAP activity in NF2 mutant meningiomas is significantly associated with time to recurrence. To this end, we combined all NF2 mutant meningiomas into one cohort (combining tumors of the aggressive and benign clusters), resulting in 464 NF2 mutant tumors with time to recurrence data. We then divided these 464 tumors into three groups of equal sample numbers based on their expression levels of a specific gene (low expression, medium expression, high expression) and compared the time to recurrence for tumors in the low expression group versus the high expression group.

**Figure 3:**
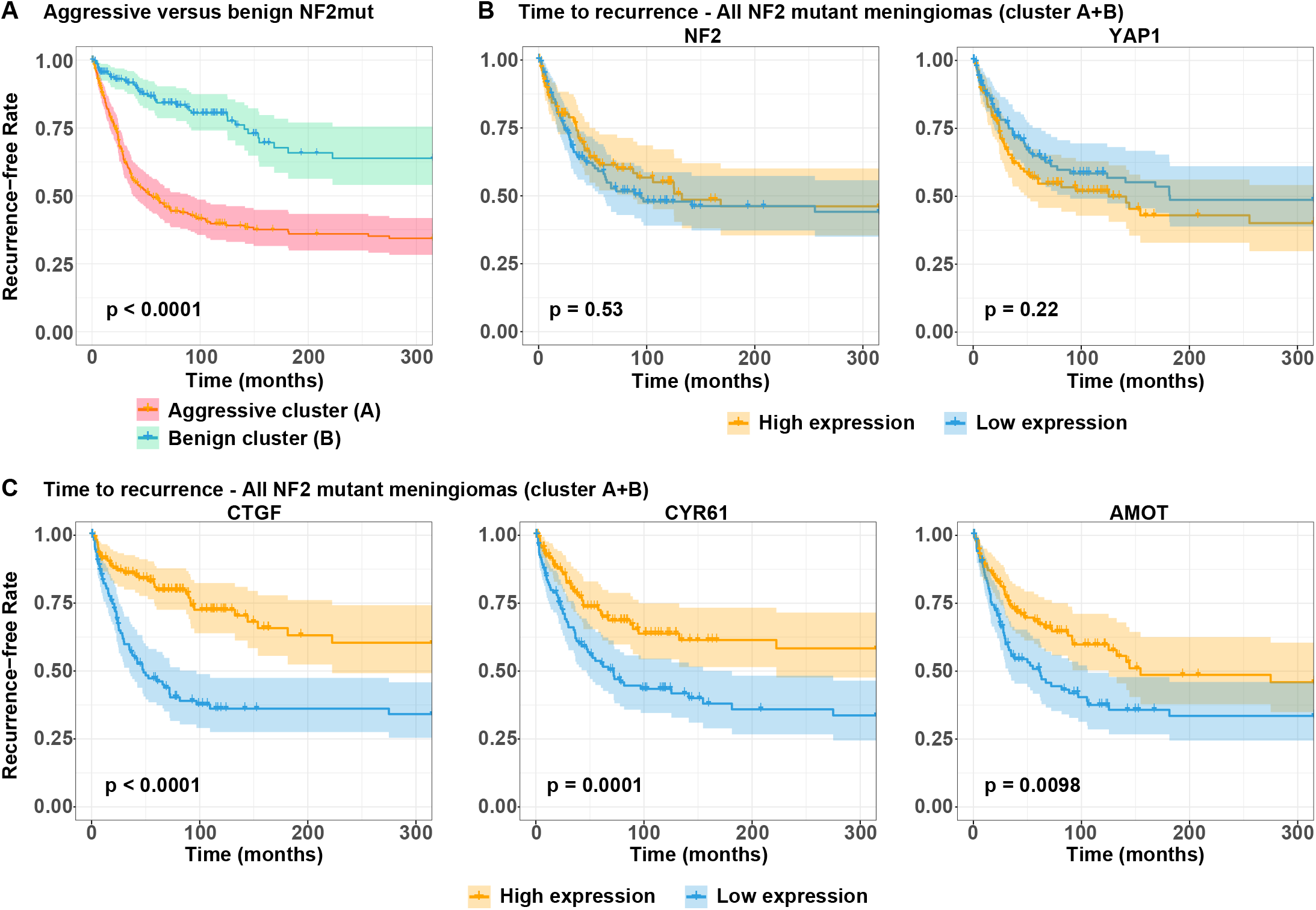
Low expression of YAP1 target genes is associated with shorter time to recurrence in NF2 mutant meningioma. A) Time to recurrence in aggressive versus benign NF2 mutant meningiomas. B-C) Time to recurrence of NF2 mutant human meningiomas harboring low or high expression of either NF2 and YAP1 (B) or the YAP targets CTGF, CYR61, and AMOT (C). Statistical analysis was done with Log-rank (Mantel-Cox) test (B, C).

We did not observe a difference in the time to recurrence for NF2 mutant meningiomas that displayed high or low expression levels of either YAP1 or NF2 (Figure 3B). By contrast, NF2 mutant meningiomas that expressed lower levels of CTGF, CYR61, or AMOT exhibited significantly shorter times to recurrence (Figure 3C). Similarly, NF2 mutant meningiomas that expressed lower levels of either CTGF, CYR61, or AMOT were enriched for tumors of higher WHO grades (Suppl. Figure S2A).

Taken together, NF2 mutant meningiomas that express lower levels of the YAP1 targets CTGF, CYR61, or AMOT are significantly associated with shorter times to recurrence and higher WHO grade.

### Aggressive NF2 mutant meningiomas up-regulate the YAP1 antagonist VGLL4 as well as the upstream inhibitors FAT3/4

In order to determine how aggressive NF2 mutant meningiomas downregulate YAP activity compared to their benign counterparts, we analyzed the expression of several regulators and mediators of YAP signaling. We analyzed the expression of different Hippo pathway upstream inhibitors of YAP1 (LATS1/2, STK3/4, MOB1A/B, SAV1, FAT1-4, FRMD6, DCHS1/2, WWC1 (KIBRA), TAOK1-3), TEAD transcription factors (TEAD1-4), as well as YAP1 competitors (VGLL1-4) in our cohort of 1298 bulk RNA-Seq samples from human meningiomas. We observed higher expression levels of several Hippo pathway members in benign NF2 mutant meningiomas, which can likely be attributed to feedback mechanisms (Suppl. Figure S3A). By contrast, we observed significantly higher expression levels of VGLL4 (2.14-fold), as well as FAT3 (10.56-fold) and FAT4 (2.27-fold) in aggressive compared to benign NF2 mutant meningiomas (Figure 4A, Suppl. Figure S3B-C). High expression of each of these three genes was associated with a significantly shorter time to recurrence in NF2 mutant tumors (Figure 4B). Importantly, high expression of both VGLL4 and FAT3 was significantly associated with shorter time to recurrence, notably within the aggressive NF2 mutant cluster itself (Suppl. Figure S3D), suggesting that these two genes are expressed at higher levels in an especially aggressive subset of tumors.

**Figure 4:**
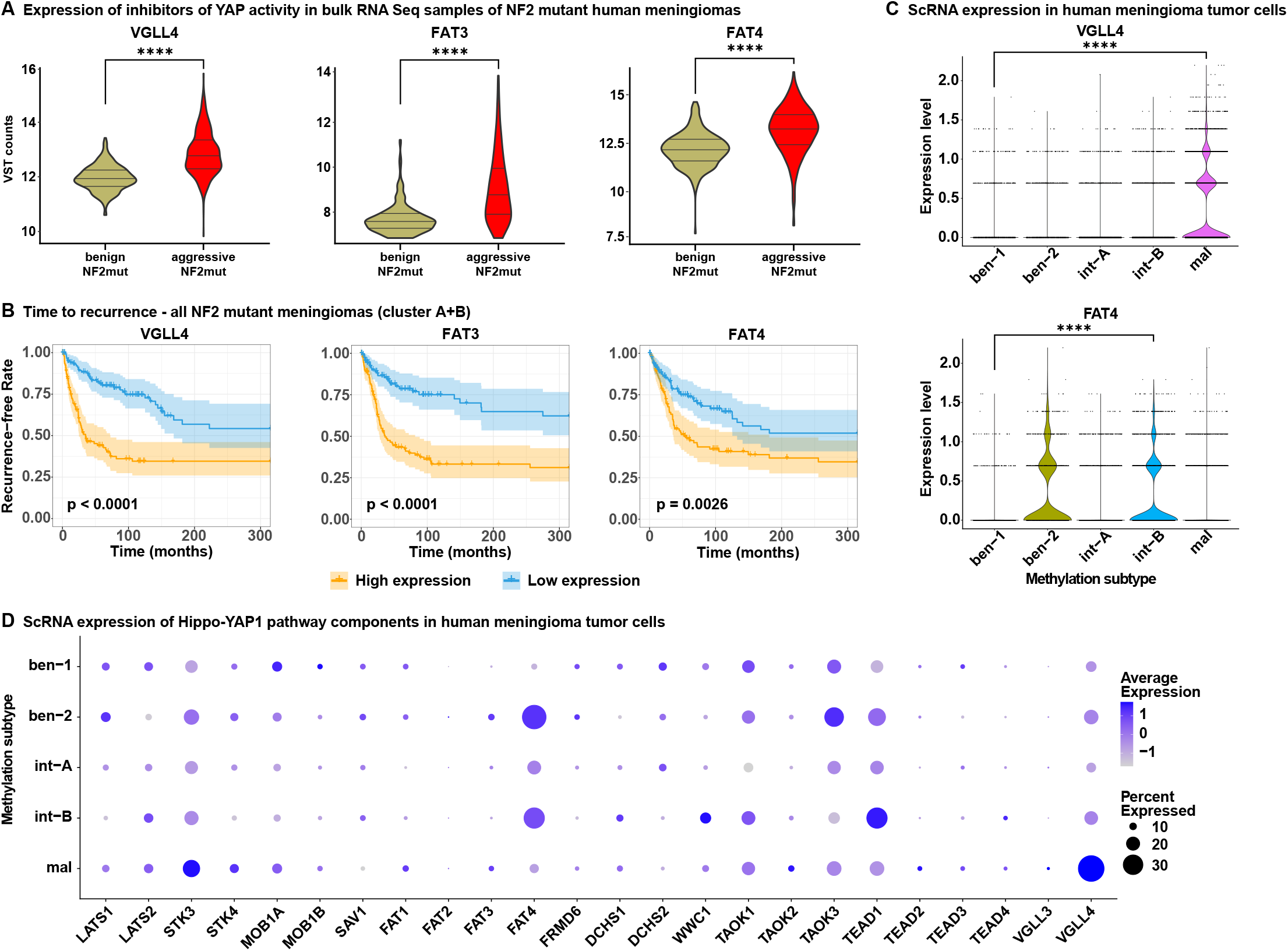
Aggressive NF2 mutant meningiomas upregulate the expression of VGLL4, FAT3, and FAT4. A) Expression of VGLL4, FAT3, and FAT4 in bulk RNA-Seq data of benign and aggressive NF2 mutant human meningiomas. B) Time to recurrence of NF2 mutant human meningiomas harboring low or high expression of either VGLL4, FAT3, or FAT4. C-D) Expression of VGLL4 and FAT4 (C) and other Hippo Pathway members (D) in single-cell RNA-Seq data of human meningioma tumor cells of different methylation subtype tumor samples. Statistical analysis was done with One-way ANOVA (B) or Log-rank (Mantel-Cox) test (A, C). (****) P _≤_ 0.0001.

We again utilized our single cell RNA sequencing data and observed increased expression levels of VGLL4 and FAT3-4 in aggressive meningioma cells. Of note, we observed significant differences in the expression of VGLL4 and FAT3-4 between different methylation-classifier subgroups. In mal-subtype tumor cells (compared to ben-1 subtype tumor cells) we observed a significantly increased expression of both VGLL4 (2.75-fold increase, padj = 4.04 x 10^-56^) as well as VGLL3, albeit at a lower absolute level (6-fold increase, padj = 0.004), compared to ben-1 tumor cells. Mal subtype cells also showed a significantly increased expression of both FAT3 (4.23-fold increase, padj = 1.13 x 10^-6^) and to a lesser degree of FAT4 (1.58-fold increase, padj = 0.018). By contrast, tumor cells of the int-B subtype did not show a significantly increased expression of VGLL4, VGLL3, or FAT3, but showed increased expression of FAT4 (4.31-fold increase, padj = 1.68 x 10^-55^) as well as WWC1 (1.99-fold increase, padj = 0.0002). Int-A subtype tumor cells, similar to int-B tumor cells, showed significantly increased expression levels of FAT4 (2.53-fold increase, p = 2.87 x 10^-17^).

Taken together, our data suggests that aggressive NF2 mutant tumors exhibit increased expression of the YAP1 antagonist VGLL4, as well as the Hippo Pathway upstream regulators FAT3 and FAT4. The mechanisms of how YAP activity is inhibited may be specific to distinct meningioma subtypes.

### Expression of the YAP1 antagonist VGLL4 leads to the inhibition of YAP activity in benign NF2 mutant meningioma cells

Since we detected increased VGLL4 expression in aggressive NF2 mutant meningioma cells, we asked what effect the overexpression of VGLL4 would have on YAP activity. To this end, we first employed a luciferase-based approach to measure the YAP activity in HEK293 cells upon VGLL4 overexpression. We transfected HEK293 cells grown at subconfluency cell densities with the YAP1 reporter plasmid (8xGTIIC-Luc), and either GFP (control), S127/397A-YAP1 (2SA-YAP1), VGLL4, as well as a combination of 2SA-YAP1 plus either GFP or VGLL4.

Overexpression of 2SA-YAP1 alone led to a significantly increased reporter activity (85.9-fold increase to GFP, p < 0.0001), whereas overexpression of VGLL4 alone resulted in significantly reduced reporter activity (0.52-fold change to GFP, p = 0.017) (Figure 5A). Furthermore, while co-expression of 2SA-YAP1 and GFP only led to a small decrease in reporter activity compared to 2SA-YAP1 alone, co-expression of VGLL4 led to a dramatic decrease in reporter activity (22.2-fold change compared to GFP). Similarly, HEK cells co-expressing 2SA-YAP1 and VGLL4 experienced significantly reduced expression levels of YAP1 target genes (CYR61, ANKRD1, AMOTL2) compared to cells expressing only 2SA-YAP1 (Figure 5B, Suppl. Figure S4A).

**Figure 5:**
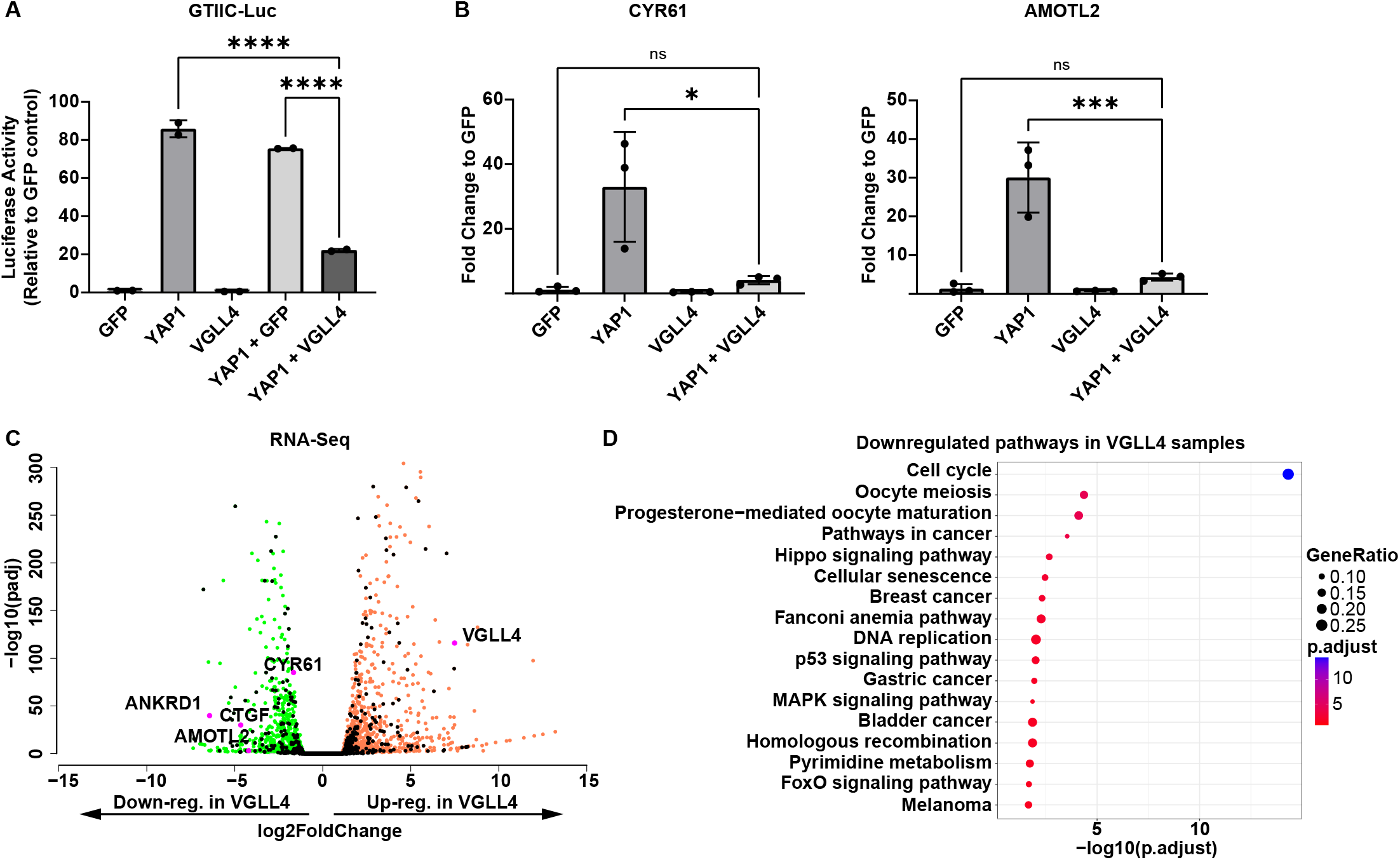
Expression of VGLL4 leads to the suppression of YAP target genes in vitro. A) Ability of VGLL4 to suppress YAP activity in the 8xGTIIC-Luc luciferase assay. B) Expression of the YAP1 target genes CYR61 and AMOTL2 in HEK cells upon transient transfection of either GFP, 2SA-YAP1, VGLL4, or 2SA-YAP1 + VGLL4. C) Overexpression of VGLL4 leads to the downregulation of YAP1 target genes in human benign NF2 mutant meningioma cells (Ben-Men-1). D) KEGG enrichment analysis of down-regulated DEGs in VGLL4-expressing Ben-Men-1 cells (compared to GFP-expressing cells). Statistical analysis was done with One-way ANOVA (A, B). (*) P _≤_ 0.05; (***) P _≤_ 0.001; (****) P _≤_ 0.0001.

To explore the effects of VGLL4 overexpression in benign NF2 mutant meningioma cells, we lentivirally transduced Ben-Men-1 meningioma cells to express either GFP or VGLL4 and performed live cell imaging as well as RNA Seq analysis. VGLL4 overexpression led to the significant de-regulation of a large number of YAP1 target genes (297 out of 2212, p = 6.8 x 10^-32^), including the downregulation of the pivotal YAP target genes CTGF, CYR61, ANKRD1 and AMOTL2 (Figure 5C). Furthermore, KEGG pathway analysis showed a significant enrichment of “Hippo Signaling pathway”-related genes in down-regulated genes in VGLL4-expressing Ben-Men-1 cells (Figure 5D).

Taken together, these results confirm the direct link between VGLL4 overexpression and reduced levels of YAP activity observed in aggressive NF2 mutant meningiomas.

## Discussion

Around 50 percent of meningiomas harbor inactivating mutations in the NF2 gene as well as loss of chromosome 22 (harboring NF2). NF2 has a myriad of tumor suppressor functions, including the regulation of the Ras-Raf-MEK-ERK and PI3K-AKT-mTOR-S6 pathways, however, one of its pivotal functions involves the negative regulation of the oncogenes YAP1 and TAZ through the Hippo Signaling pathway. Loss of NF2 results in deregulation and activation of oncogenic YAP activity and increased levels of YAP activity have been observed in NF2 mutant over NF2 wild type meningiomas ^11,12^. Oncogenic YAP activity has also been implicated in the pathobiology of other tumors harboring inactivating NF2 mutations ^44-47^. We have previously shown that the expression of an oncogenic point mutant YAP construct is able to induce the formation of meningioma-like tumors in mice ^13^. This link between NF2 and YAP is further highlighted by the presence of YAP1 gene fusions in pediatric NF2 wild type meningiomas, which cluster closely with NF2 mutant meningiomas based on DNA methylation and RNA Seq classifications ^13,14^.

While inactivating NF2 mutations and loss of chromosome 22 are frequently the only recurrent mutations in benign NF2 mutant meningiomas, higher-grade and aggressive NF2 mutant tumors generally harbor a more aberrant genome, suggesting that additional pathways are de-regulated in these tumors. Our data suggests that aggressive NF2 mutant meningiomas downregulate at least a subset of the oncogenic YAP1 activity that is present in benign NF2 mutant meningiomas. We observed this effect across tumors of all higher-grade methylation subtypes (int-A, int-B, mal) when compared to tumors of the benign ben-1 subtype. Our data furthermore indicates that this down-regulation of YAP activity can be achieved in several ways. Tumors of the mal subtype significantly upregulated the expression of the transcriptional co-factor VGLL4 (and to a lesser degree VGLL3). VGLL proteins, similar to YAP1, bind to TEAD transcription factors, thereby preventing the interaction between YAP1 and TEADs. Tumors of all aggressive subtypes, while more pronounced in tumors of the int-A and int-B subtypes, also overexpressed FAT4. Mal subtype tumors also upregulated the expression of FAT3. In Drosophila, FAT proteins have been shown to activate the Hippo Signaling pathway and suppress YAP activity independently of NF2 function. It cannot be excluded that additional, yet unknown mechanisms contribute to the attenuation of YAP activity in aggressive NF2 mutant meningiomas.

In vitro data with Ben-Men-1 benign NF2 mutant meningioma cells and HEK293 cells showed that overexpression of VGLL4 results in the significant suppression of YAP1 activity, confirming the direct link between overexpression of VGLL4 and reduced YAP activity. Functional inhibition or loss of expression of YAP1 has been observed in several cancers, such as Estrogen receptor (ER)-positive breast cancer, neuroendocrine small-cell lung cancer (SCLC), or pulmonary large cell neuroendocrine carcinoma ^48-51^. YAP1 was originally identified as a tumor suppressor due to its role in activating p73-dependent apoptosis ^52,53^ and has been shown to suppress ER expression in ER+ breast cancer as well as suppress metastasis in SCLC ^49,51^. However, the reasons why YAP activity is inhibited in aggressive NF2 mutant meningiomas remain unclear and warrant further investigation.

It is currently unknown if benign and aggressive NF2 mutant meningiomas equally rely on YAP1 signaling for their growth and/or survival. YAP1 is a transcriptional co-activator that does not bind DNA itself, but instead relies on the interaction with other transcription factors (mostly TEAD1-4). Several pharmacological inhibitors of YAP signaling are currently being developed, most of them inhibiting the interaction of YAP1 with TEAD transcription factors, and their efficacy is currently being assessed in phase 1 clinical trials of several YAP1-activated tumor types (such as NF2 mutant mesothelioma) ^24,54^. It remains unclear if these inhibitors are equally effective against both benign (that seem to largely rely on YAP signaling) and aggressive NF2 mutant meningiomas (that harbor additional mutations and likely concurrently activate additional mitogenic pathways), especially since aggressive tumors show decreased baseline levels of YAP activity.

In summary, our study suggests that aggressive NF2 mutant tumors downregulate oncogenic YAP activity, in part by upregulating the expression of the YAP1 antagonist VGLL4 and the upstream regulators FAT3/4. Our findings may have important implications for the efficacy of therapeutic approaches aimed at the YAP1-TEAD complex in benign versus aggressive NF2 mutant meningiomas.

## Material & Methods

### RNA sequencing data of human meningiomas

The collection of bulk RNA Seq data from human meningiomas has previously been published ^41^. Single-cell RNA Seq data of human meningioma tumors has previously been published ^43^, was obtained from the data repository of the Dept. of Neuropathology at the University Hospital Heidelberg and is available upon request. RNA-Seq data from human Ben-Men-1 cells expressing either VGLL4 or GFP can be accessed at the GEO database at GSE263122. The code used to process and analyze the data is available at https://github.com/sonali-bioc/SzulzewskyVGLL4Paper.

### Bulk RNA-Seq analysis

For analysis of human meningioma tumors, Bulk RNASeq data (vst counts), UMAP coordinates and metadata for 1298 human Meningioma Tumors from ^41^ was downloaded. R package DESeq2 was used to find differentially expressed genes (DEGs) between Cluster A (NF2 mutant aggressive) vs Cluster B (NF2 mutant benign). A threshold of fold change of 1.5 (or logFC of 0.58) and a p-adjusted value < 0.05 was used to determine significantly regulated DEGs. For analysis of RNA-Seq data from Ben-Men-1 cells expressing either VGLL4 or GFP, Raw sequencing reads were checked for quality using FastQC (https://www.bioinformatics.babraham.ac.uk/projects/fastqc/). RNA-seq reads were aligned to the hg38 assembly using gencode v39 and STAR2 ^55^ and counted for gene associations against the UCSC genes database with HTSeq ^56^. Differential Expression analysis for RNASeq Data was performed using R/Bioconductor package DESeq2 ^57^ and edgeR ^58^. A log2fold change cutoff of 1 (fold change of 2) and FDR < 0.05 was used to find transcriptionally regulated genes.

### Survival Analysis

For each gene, using the gene expression for samples present in cluster A and B, the patient samples were divided into 2 groups – group1 contained samples whose gene expression was less than 1st quantile and group2 contained samples whose gene expression was higher than 3rd quantile of all samples. R package survival (https://cran.r-project.org/web/packages/survival/index.html) and survminer (
https://cran.r-project.org/web/packages/survminer/index.html), was then used to draw a KM curve and find survival differences between the two groups.

### Single-cell RNA-Seq data analysis

Seurat objects were constructed for 26 single-cell RNA-Seq samples of human meningiomas ^59^. Cells from the 26 single-cell RNA-Seq samples were further filtered to remove immune cells based on 49 marker genes (Suppl. Table S1) spanning a variety of immune cell types. Next SCTransform from Seurat was used to process the data, followed by building a UMAP and forming clusters. Metadata extracted for the remainder cells was used to divide cells into different groups for WHO and methylation cluster (MC) status. Dotplots and violin Plots for key genes were made using the function DotPlot() and VlnPlot() respectively from Seurat. FindMarkers() from Seurat was used to find differentially expressed genes between the different MC status (ben-1 vs ben-2, ben-1 vs int-A, ben-1 vs int-B, ben-1 vs mal). R package, R function, DoHeatmap() was used to make heatmaps. All analysis was done using R 4.3.3 and Seurat (v 5.0.0).

See also Suppl. Methods.

## Author Contributions

Conceptualization ECH, FS. Performed experiments AP, DR, SS, FS. Data analysis AP, SA, HNT, PS, FS. Original manuscript writing ECH, FS. Review and editing SA, ECH, FS. Funding acquisition ECH. Supervision FSahm, ECH, FS. All authors read, reviewed, and approved the manuscript.

## Acknowledgements

We thank Denis Adair, Linda Lew, Kelly Grissom, James Yan, and Debra K Kumasaka for continued administrative assistance and support throughout these experiments. We thank Alyssa Dawson, Elizabeth Jensen, and Dolores Covarrubias at the Fred Hutchinson Genomics Core for help with DNA sequencing.

## Data and materials availability

The data that support the findings of this study are included with the manuscript and supplemental data files and are also available from the corresponding author upon reasonable request.

## Supplementary Information

Supplementary Figures S1-S4

Supplementary Figure Legends

Supplementary Methods

Supplementary Table S1

Supplementary Table S2

## Supplementary Table S1

A) Sample info of the 26 single-cell RNA-Seq samples of human meningiomas. B) List of marker genes used to filter out immune cells from single-cell RNA-Seq data.

## Supplementary Table S2

A) List of primers used/ B) List of plasmids used.

