## Supplemental Data for "Aggressive high-grade NF2 mutant meningiomas downregulate oncogenic YAP signaling via the upregulation of VGLL4 and FAT3/4"

Supplementary Information

Table of Contents:

Supplementary Figures S1-S4
Supplementary Figure Legends
Supplementary Methods
Supplementary References

**Suppl. Figure S1**

### A Baylor RNA classification

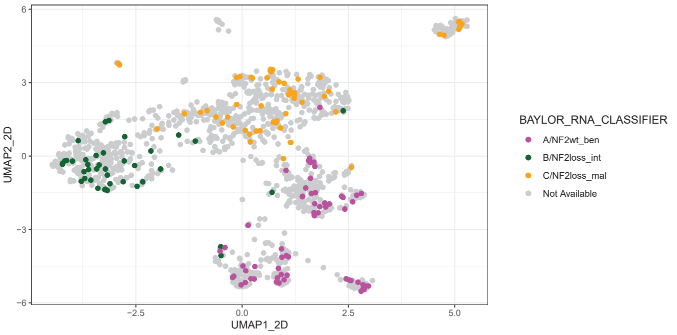


### B Heidelberg DNA methylation classification

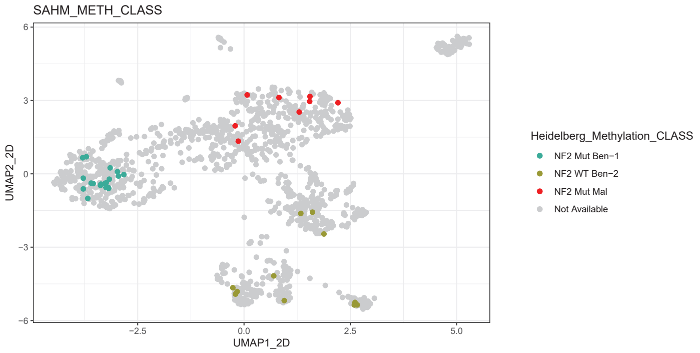


### C Expression of NF2, YAP1 and YAP1 targets in human meningiomas

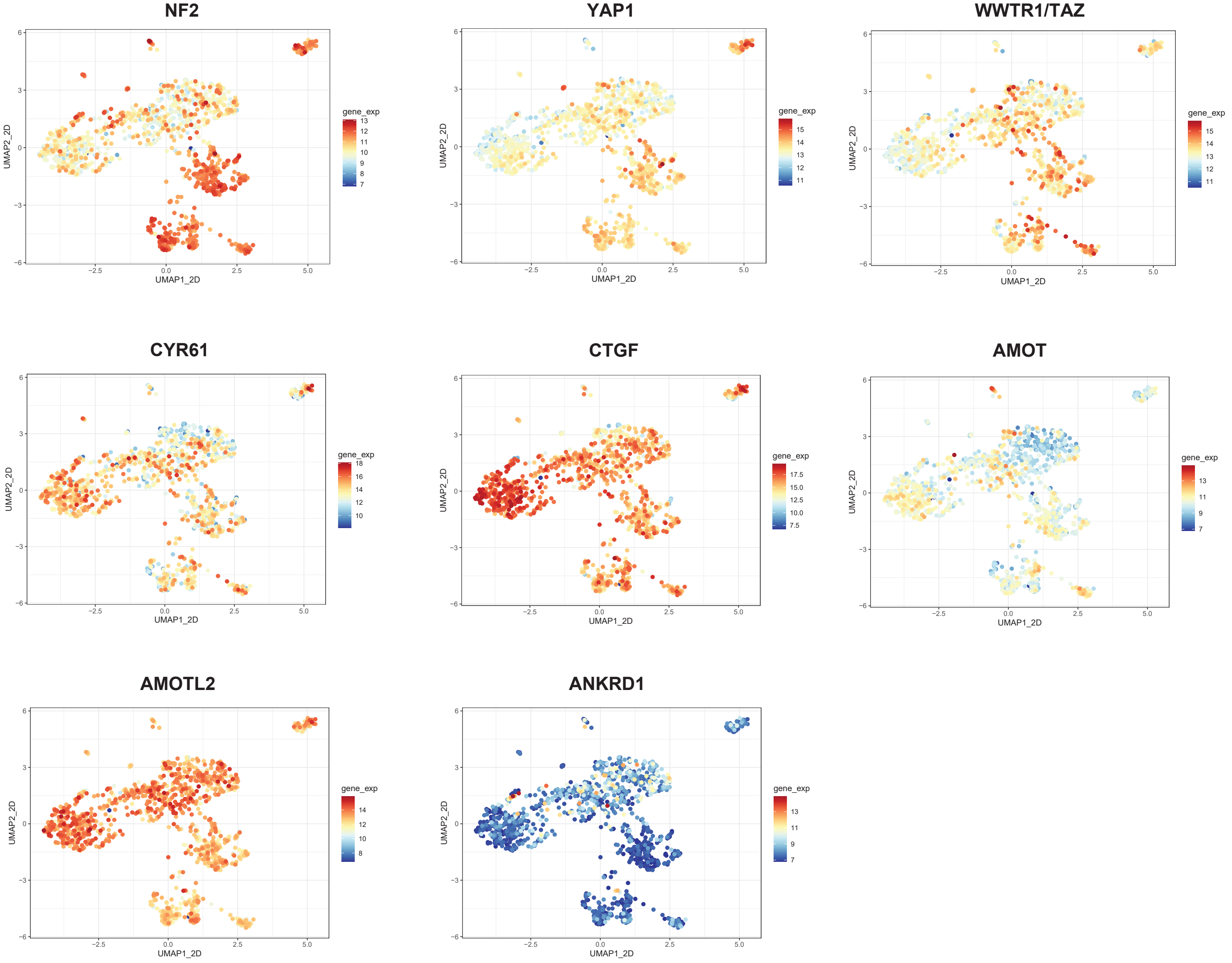

### Suppl. Figure S2

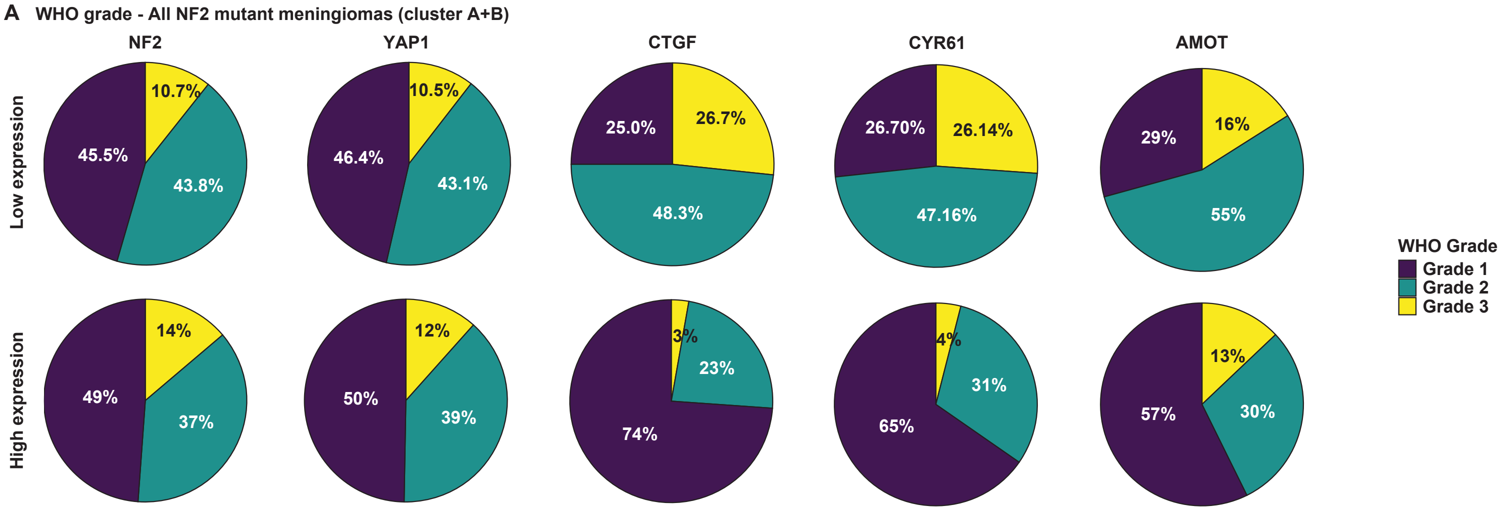

### Suppl. Figure S3A

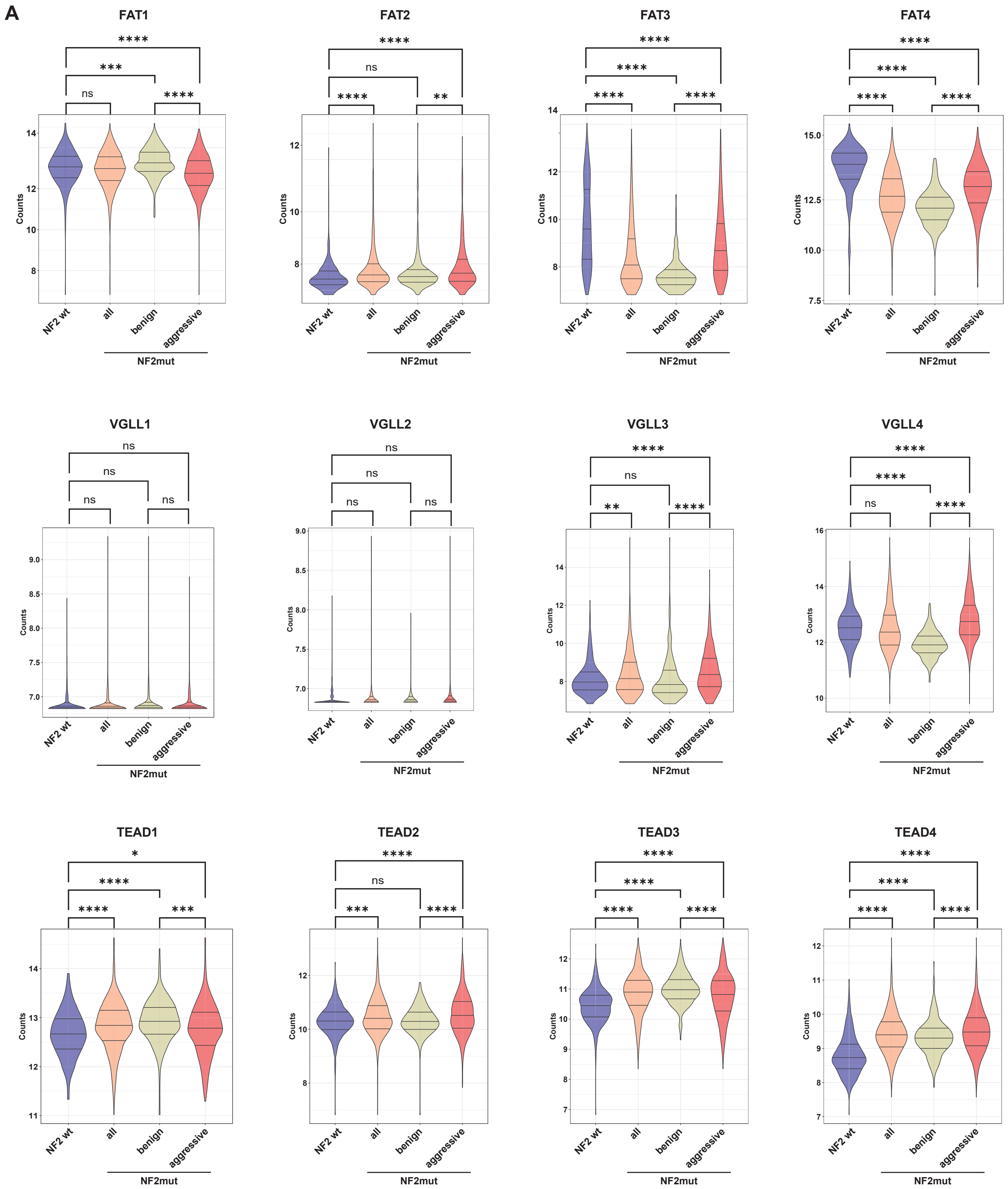

### Suppl. Figure S3A (continued)

### A (continued)

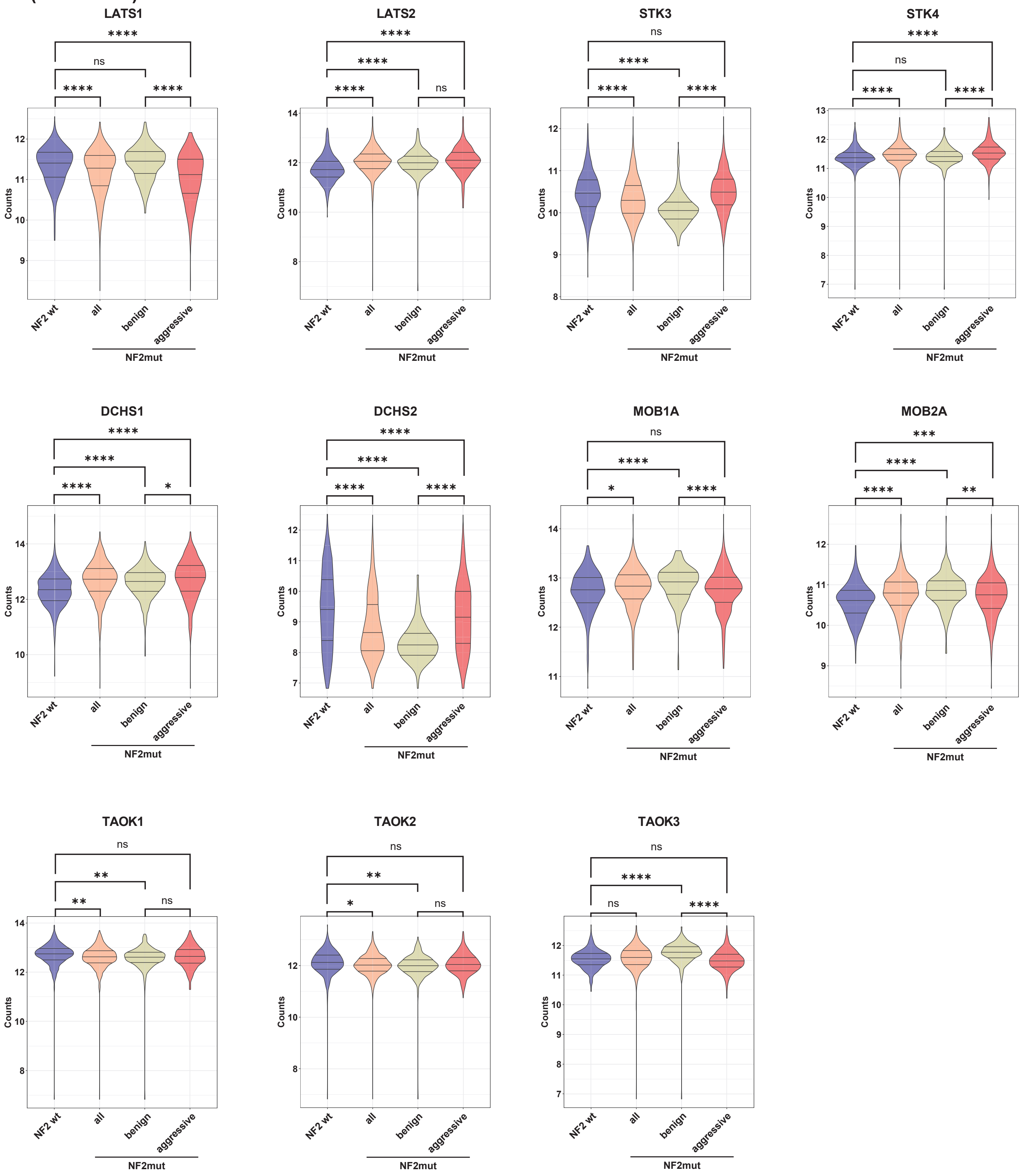

### Suppl. Figure S3A (continued)-C

### A (continued)

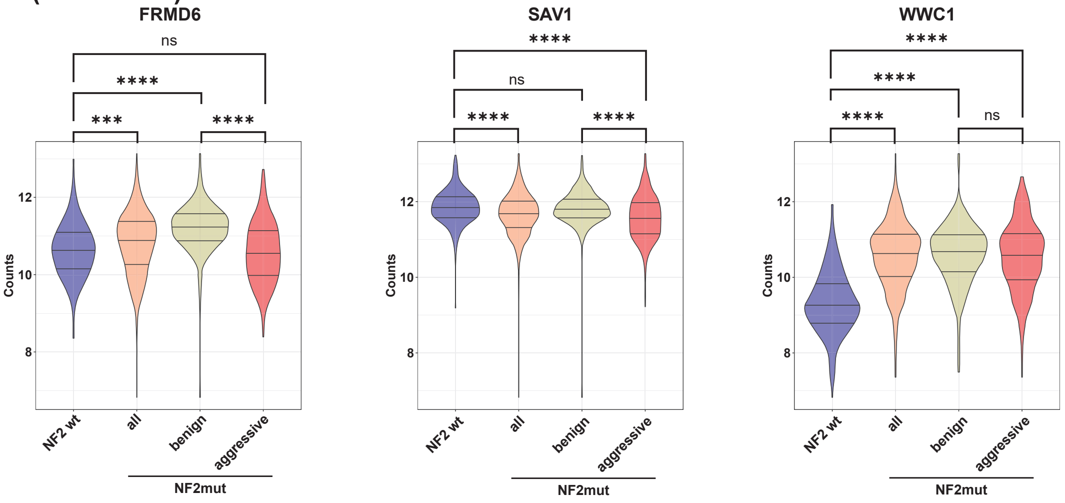


## B

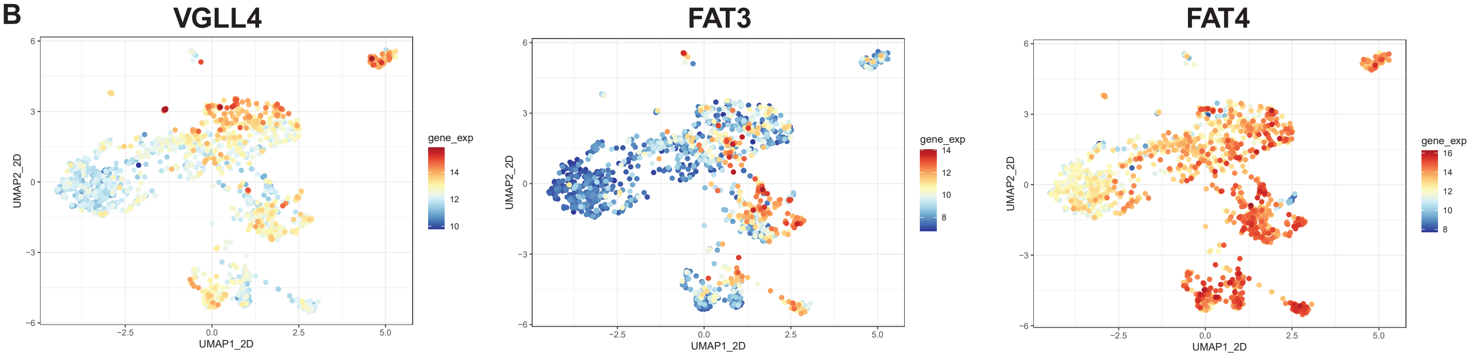


### C Time to recurrence - all NF2 mutant meningiomas (cluster A+B)

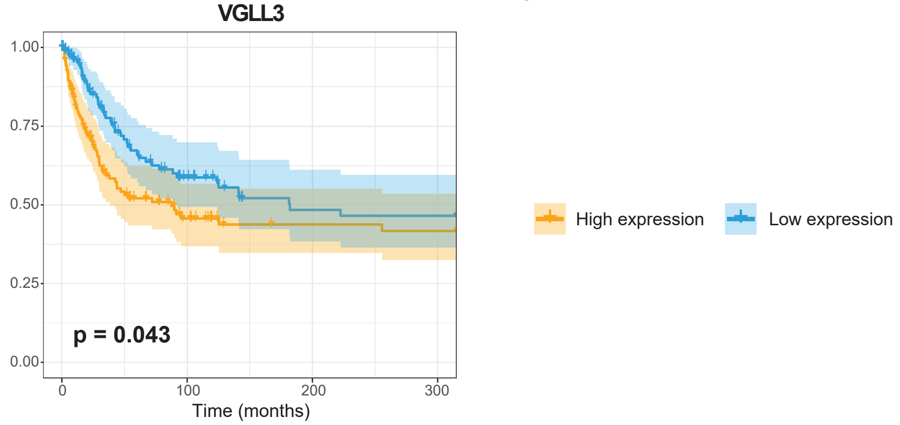

### Suppl. Figure S3D

**D** **Time to recurrence - only NF2 mutant meningiomas in the aggressive cluster (cluster A)**

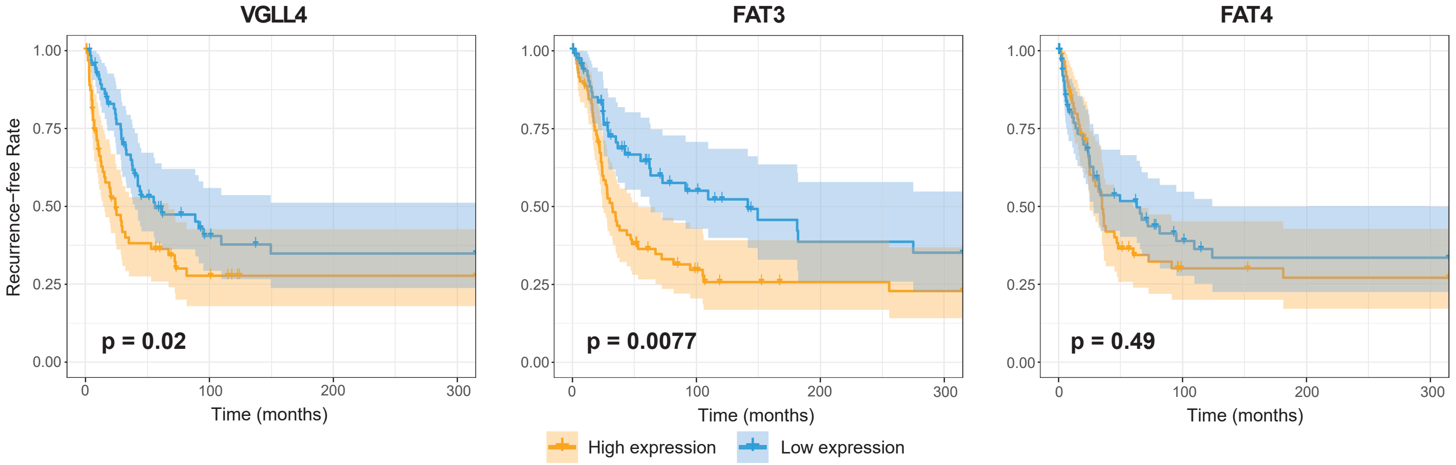

### Suppl. Figure S4

A

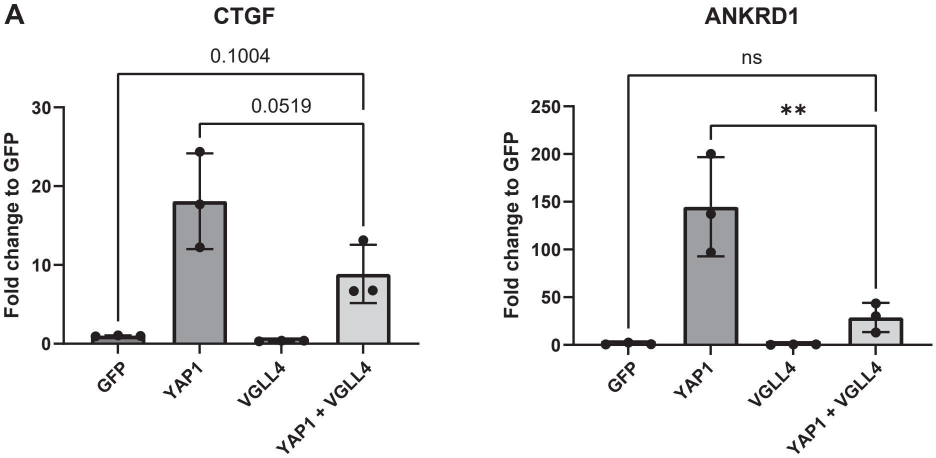

**Suppl. Figure legends:**

**Suppl. Figure S1: Aggressive NF2 mutant meningiomas display decreased levels of YAP activity.** A-C) Reference UMAPs showing clustering of human meningiomas based on bulk RNA-Seq data. Samples are colored by Baylor RNA classification status (A), Heidelberg methylation classifier status (B), or gene expression of several Hippo Pathway members and YAP1 target genes (C). Statistical analysis was done with One-way ANOVA (B). (***) $P \leq 0.001$; (****) $P \leq 0.0001$.

**Suppl. Figure S2: Low expression of YAP1 target genes is associated with shorter time to recurrence in NF2 mutant meningioma.** A) WHO grades of tumors harboring low or high expression of either NF2 and YAP1 or the YAP targets CTGF, CYR61, and AMOT.

**Suppl. Figure S3: Aggressive NF2 mutant meningiomas upregulate the expression of VGLL4, FAT3, and FAT4.** A) Expression (VST counts) of Hippo Pathway genes in bulk RNA-Seq data of NF2 wild type, as well as NF2 mutant (benign and aggressive) human meningiomas. B) Reference UMAPs showing clustering of human meningiomas based on bulk RNA-Seq data. Samples are colored by gene expression of VGLL4, FAT3, and FAT4. C) Time to recurrence of NF2 mutant human meningiomas tumors harboring low or high expression of VGLL3. D) Time to recurrence of NF2 mutant human meningiomas (aggressive subtype tumors only) harboring low or high expression of VGLL4, FAT3, or FAT4. Statistical analysis was done with One-way ANOVA (A) or Log-rank (Mantel-Cox) test (C, D). (*) $P \leq 0.05$; (**) $P \leq 0.01$; (***) $P \leq 0.001$; (****) $P \leq 0.0001$.

**Suppl. Figure S4: Expression of VGLL4 leads to the suppression of YAP target genes in vitro.** A) Expression of the YAP1 target genes CTGF and ANKRD1 in HEK cells upon transient transfection of either GFP, 2SA-YAP1, VGLL4, or 2SA-YAP1 + VGLL4. Statistical analysis was done with One-way ANOVA (A). (**) $P \leq 0.01$.

**Suppl. Methods**

**Enrichment Analysis and Data visualization**
Gene Set Enrichment Analysis (GSEA) was performed against the MsigDB database with the KEGG gene sets using enrichR (1) package. Resulting enriched genesets and pathways were filtered via a threshold of FDR < 0.05. Fisher hypergeometric tests were implemented in R using function phyper() to see if genes in one set were over-represented, compared to other gene sets. R package pheatmap(https://cran.r-project.org/web/packages/pheatmap/index.html ) was used to make heatmaps. Volcano plots were made using R (v4.3.3). All other plots were made using ggplot2.

**Plasmid preparation**
Primers used for plasmid generation are listed in Suppl. Table S2A. Additional plasmids used in this study are listed in Suppl. Table S2B.

**Luciferase Assays**
HEK293 cells were cultured in DMEM, 10% FBS, 1% Penicillin/Streptomycin. If not indicated differently, HEK293 cells were seeded into white 96 well plates at 10,000 cells/well the day prior to transfection. Cells were then transfected with the indicated plasmids and a plasmid containing Renilla using Lipofectamin 3000 (Thermo Fisher Scientific) according to the manufacturer's instructions. Luciferase activity was measured 24 hours after transfection using the Dual-Glo Luciferase Assay System (Promega) on a Veritas Microplate Luminometer.

**Lentiviral transductions**
For virus production, pLJM1 (Addgene) constructs containing the inserts of interest were transfected into 293T cells, along with psPAX and pMD2.G packaging plasmids (Addgene), using polyethylenimine (Polysciences). Fresh media was added 24 hours later, and viral supernatant harvested 24 hours after that. For infection of NIH3T3 cells, 1x105 cells/well were seeded into 6-well plates. Lentivirus was used unconcentrated and cells were infected at a MOI<1 24 hours after seeding. 72 hours after seeding, selection was begun for cells successfully expressing the constructs using 1 µg/mL puromycin (for 3 days).

**RNA isolation, PCR, and RNA Sequencing**
RNA was extracted using the Qiagen RNeasy Mini Kit according to the manufacturer's instructions. Genomic DNA was removed by on-column DNase digestion. Total RNA integrity was checked using an Agilent 4200 TapeStation (Agilent Technologies, Inc., Santa Clara, CA) and quantified using a Trinean DropSense96 spectrophotometer (Caliper Life Sciences, Hopkinton, MA). RNA-seq libraries were prepared from total RNA using the TruSeq Stranded mRNA kit (Illumina, Inc., San Diego, CA, USA). Library size distribution was validated using an Agilent 4200 TapeStation (Agilent Technologies, Santa Clara, CA, USA). Additional library QC, blending of pooled indexed libraries, and cluster optimization was performed using Life Technologies' Invitrogen Qubit® 2.0 Fluorometer (Life Technologies-Invitrogen, Carlsbad, CA, USA). RNA-seq libraries were pooled (70-plex) and clustered onto a P3 flow cell. Sequencing was performed using an Illumina NextSeq 2000 employing a paired-end, 50 base read length (PE50) sequencing strategy. For quantitative real-time PCR, RNA was transcribed into cDNA using the SuperScript III kit. PCR experiments were carried out on a QuantStudio™ 7 Flex Real-Time PCR System. For primer sequences see Suppl. Table S2A.

### Suppl. References

1. Chen EY, Tan CM, Kou Y, Duan Q, Wang Z, Meirelles GV, et al. Enrichr: interactive and collaborative HTML5 gene list enrichment analysis tool. BMC Bioinformatics. 2013;14:128.
